# Targeting Age-Related Differences in Brain and Cognition with Multimodal Imaging and Connectome Topography Profiling

**DOI:** 10.1101/601146

**Authors:** Alexander J. Lowe, Casey Paquola, Reinder Vos de Wael, Manesh Girn, Sara Lariviere, Shahin Tavakol, Benoit Caldairou, Jessica Royer, Dewi V. Schrader, Andrea Bernasconi, Neda Bernasconi, R. Nathan Spreng, Boris C. Bernhardt

**Author notes:** CORRESPONDING AUTHOR, Boris Bernhardt, PhD, Multimodal Imaging and Connectome Analysis Lab, McConnell Brain Imaging Centre, Montreal Neurological Institute and Hospital, McGill University, Montreal, Quebec, H3A 2B4, Canada.

## Abstract

Aging is characterised by accumulation of structural and metabolic changes in the brain. Recent studies suggest transmodal brain networks are especially sensitive to aging, which, we hypothesise, may be due to their apical position in the cortical hierarchy. Studying an open-access healthy cohort (n=102, age range = 30-89 years) with MRI and Aβ PET data, we estimated age-related cortical thinning, hippocampal atrophy and Aβ deposition. In addition to carrying out surface-based morphological and metabolic mapping, we stratified effects along neocortical and hippocampal resting-state functional connectome gradients derived from independent datasets. The cortical gradient depicts an axis of functional differentiation from sensory-motor regions to transmodal regions, whereas the hippocampal gradient recapitulates its long-axis. While age-related thinning and increased Aβ deposition occurred across the entire cortical topography, increased Aβ deposition was especially pronounced towards higher-order transmodal regions. Age-related atrophy was greater towards the posterior end of the hippocampal long-axis. No significant effect of age on Aβ deposition in the hippocampus was observed. Imaging markers correlated with behavioural measures of fluid intelligence and episodic memory in a topography-specific manner. Our results strengthen existing evidence of structural and metabolic change in the aging brain and support the use of connectivity gradients as a compact framework to analyse and conceptualize brain-based biomarkers of aging.

## I. Introduction

Aging is a multifactorial process defined as a time-dependant functional decline that affects most living organisms (López-Otín et al., 2013). Though not fully understood, aging involves the accumulation of structural and metabolic changes that ultimately lead to impairments in multiple cognitive domains, including executive function, episodic memory, and word retrieval (Baciu et al., 2016; Fjell et al., 2017; Tromp et al., 2015). Collectively, these contribute to increasing challenges for psychosocial functioning, wellbeing, and quality of life (Pan et al., 2015; Wilson et al., 2013).

Ongoing advances in multimodal neuroimaging have identified structural, functional, and metabolic substrates of both healthy cognitive functioning (Tomasi and Volkow, 2012) and of its decline in aging (Draganski et al., 2013; McConathy & Sheline, 2015; Steffener et al., 2013). Progress in Magnetic Resonance Imaging (MRI) acquisition and modelling techniques now allows for the fine-grained mapping of neocortical and subcortical morphology *in-vivo* (B. Fischl & Dale, 2000). In healthy individuals, age is associated with decreased hippocampal volume and cortical thinning (Fjell et al., 2014; Fraser et al., 2015; Salat et al., 2004; Shaw et al., 2016; Sowell et al., 2003; Yao et al., 2012; Yang et al., 2016), both of which measurably contribute to cognitive decline (Fjell et al., 2006; Leal & Yassa, 2015; Mielke et al., 2012; Walhovd et al., 2006). These findings are complemented by functional and metabolic studies, particularly work based on positron emission imaging (PET) of tracers sensitive to deposits associated with healthy and pathological aging. Notably, PET-based quantification of amyloid beta (Aβ), generally considered a marker of neurodegenerative conditions like Alzheimer’s Disease, has demonstrated elevated levels in the brains of cognitively normal older adults as well (Jansen et al., 2015; Rodrigue, Kennedy, & Devous, 2012; Sperling et al., 2011). Cortical Aβ has furthermore been associated with multi-domain cognitive impairment, such as episodic and semantic memory together with executive and visuospatial abilities (Baker et al., 2017; Farrell et al., 2017; Jansen et al., 2018; Mortamais et al., 2017).

With increasing availability of open-access and multimodal data aggregation and dissemination initiatives, it is now possible to adopt an integrated approach, combining several imaging markers to better understand biological factors contributing to cognitive decline. Recent studies in cognitively normal older adults have suggested a synergistic relationship between cerebral amyloid pathology and hippocampal atrophy (Bilgel et al., 2018), whereas others suggest that cortical thickness may represent a more approximate marker of the pathophysiological underpinning of cognitive decline than Aβ deposition (Knopman et al., 2018). In addition to the potentially complementary value of imaging markers as surrogates of cognitive abilities, aging effects do not seem to be uniform across different regions (McGinnis et al., 2011). Notably, several reports have supported a role of large-scale functional network topology, and changes in structural covariance networks, on the risk for cognitive decline in aging (Andrews-Hanna et al., 2007; Buckner, 2004; Fox et al., 2005; Sambataro et al., 2010; Spreng & Turner, 2013; Spreng et al., 2010; Zhao et al., 2015). Structural and metabolic changes in transmodal regions known to engage in more higher-order and integrative processing, such as the default mode network (DMN) and frontoparietal networks, have been demonstrated to contribute to cognitive decline in healthy subjects (Lim et al., 2014). An overlap between elements of the DMN and Aβ pathology has been previously reported in Alzheimer’s Disease (Buckner et al., 2005), with recent studies indicating that core DMN regions being among the earliest sites of Aβ deposition in preclinical stages (Palmqvist et al., 2017). This pathological accumulation is thought to contribute to memory dysfunction associated with dementia, as the DMN has been shown to be engaged during activation of episodic memory (Buckner et al., 2008). Pathological Aβ deposition is not unique to the DMN however, with early accumulation also reported in the fronto-parietal network as well as other transmodal regions with high connectivity (Buckner et al., 2009; Elman et al., 2016; He et al., 2014; Wang et al., 2007).

Studying the openly-available Dallas Lifespan Aging Study (DLBS) dataset (Park, 2018), the current work integrated measures of neocortical and hippocampal morphology and Aβ deposition to examine age-related differences and their relationship to cognition. In addition to leveraging surface-based processing and multimodal co-registration techniques, we harnessed a novel analysis reference frame determined by the putative neocortical hierarchy. Initially formulated in non-human primates (Mesulam, 1998), the hierarchy follows a ‘sensory-fugal’ gradient from low-level cortices involved in interactions with the external world to higher-order transmodal areas involved in self-generated, abstract cognition (Buckner and Krienen, 2013; Huntenburg et al., 2018; Margulies et al., 2016; Paquola et al., 2018). Recent application of unsupervised compression techniques applied to cortico-cortical functional connectivity data recapitulated a similar gradient in humans (Margulies et al., 2016). By being functionally and anatomically distant from sensory systems, DMN activity is likely to be shielded from environmental input (Kiebel, Daunizeau, & Friston, 2008) and may also perform cross-modal integration of information (van den Heuvel & Sporns, 2013). Equivalent compression techniques have been applied to hippocampus-to-cortex connectivity profiles, also revealing a principal gradient of connectivity. In the hippocampus, this gradient follows its ‘long axis’, with anterior segments being closely connected to transmodal DMN and temporo-limbic networks, while posterior sections increasingly interact with posterior cortical areas including the visual and dorsal/ventral attention networks (Vos de Wael et al., 2018).

Stratification of aging biomarkers based on neocortical and hippocampal connectivity gradients, provides an alternative viewpoint to voxel-or parcellation-based studies of macroscale organization and connectivity which in turn, may complement recent literature demonstrating increased vulnerability of higher-order networks to pathological protein accumulation (Buckner et al., 2009; Elman et al., 2016; He et al., 2014; Palmqvist et al., 2017), and age-related reductions in cortical thickness across higher-order and sensorimotor networks (Bajaj et al., 2017). Furthermore, by sidestepping the need to define discrete communities through the use of clustering (Eickhoff, Yeo, & Genon, 2018; Yeo et al., 2011) or connectivity boundary mapping techniques (Cohen et al., 2008), topographic connectome profiling provides a continuous coordinate system to aggregate and analyze aging biomarkers, allowing for the study of pathological advance along a quantifiable map of the neocortical hierarchy and hippocampal long-axis, ultimately furthering our understanding of neurological aging and the associated cognitive decline.

## II. Materials and Methods

### Participants

We selected 144 healthy adult native English speakers (89 females, 30-89 years, mean±SD age= 62±16.9 years, 93.4% White/Caucasian) from the openly-shared DLBS, a comprehensive study designed to understand cognitive functioning at different stages of the adult lifespan [http://fcon_1000.projects.nitrc.org/indi/retro/dlbs.html; (Park, 2018)]. Participants were well educated (mean±SD=16.8±2.3 years of education) and scored highly on the Mini-Mental State Examination [MMSE; (Folstein et al., 1975); mean±SD=28±1.2 points]. As previously outlined (Rodrigue et al., 2012), participants were screened for neurological and psychiatric disorders, loss of consciousness >10 minutes from a traumatic insult to the head, drug/alcohol abuse, stroke and major heart surgery, chemotherapy within 5 years, and immune system disorders. Participants were right-handed and recruited from the Dallas-Fort Worth metropolitan area. Specifically, we selected only those who underwent a research-dedicated anatomical MRI and Aβ PET examination.

The DLBS was approved by the University of Texas at Dallas and University of Texas Southwestern Medical Centre respective ethics committees. All DLBS subjects provided written consent prior to enrolment.

### Neuropsychological Test Battery

Participants completed a battery of neuropsychological tests assessing the following domains; processing speed (Salthouse & Babcock, 1991; Wechsler, 2008), working memory (Turner & Engle, 1989; Wechsler, 2008), episodic memory (Brandt, 1991; Robbins et al., 2010), crystallized abilities (Zachary, 1986), executive function (Robbins et al., 2010), and fluid reasoning (Ekstrom et al., 1976; Raven, 1996). A single subject was missing a single score for the Digit Symbol Task (0.1% missing data). Due to this number being so small, we did not want to exclude the subject from analysis, but instead imputed the missing data point using linear interpolation. Results from all neuropsychological tests were standardized to z-scores. To reduce data dimensionality, we iteratively performed a maximum likelihood common factor analysis with varimax rotation with two-to five-factor solutions (Harman, 1976). Overfitting occurred with three or more factors (Heywood, 1931), thus the two-factor solution was utilised in subsequent analyses. Neuropsychological tests pertaining to fluid intelligence strongly contributed to the first factor (*henceforth*, F1), specifically Ravens Progressive matrices, Educational Testing Service (ETS) Letter Sets, Digit Symbol Test, Digit Comparison Test, CANTAB Spatial Working Memory Test and CANTAB Stockings of Cambridge Test. Whereas tests pertaining to episodic memory contributed highly to the second factor 2 (F2), specifically Hopkins Verbal Learning Immediate Recall, Delayed Recall, and Recognition. For specific factor loadings across tests, see **Supplementary Table 1**.

### MRI Acquisition

Anatomical images were acquired with a Philips Achieva 3T whole-body scanner (Philips Medical Systems, Bothell, WA) and a Philips 8-channel head coil at the University of Texas Southwestern Medical Center using the Philips SENSE parallel acquisition technique. A 3D T1-weighted sagittal magnetization-prepared rapid acquisition gradient echo (MPRAGE) structural image was obtained (T1w, Repetition time [TR]=8.1 ms, echo time [TE]=3.7 ms, flip-angle=12°, FOV=204×256 mm^2^, resulting in 160 slices with 1×1×1 mm^3^ voxels).

### Amyloid PET Acquisition

All participants were injected with a 370 MBq (10mCi) bolus of ^18^F-Florbetapir. At 30 minutes post-injection, subjects were positioned on the imaging table of a Siemens ECAT HR PET scanner. Velcro straps and foam wedges were used to secure the participants head and participant positioning was completed using laser guides. A 2-min scout was acquired to ensure the brain was in the field of view and that there was no rotation in either plane. At 50 minutes post injection, a 2-frame by 5-minute dynamic emission acquisition was started, followed immediately by a 7-minute internal rod source transmission scan. The transmission image was reconstructed using backprojection and a 6-mm full-width-at-half-maximum (FWHM) Gaussian filter. Emission images were processed by iterative re-construction, specifically 4 iterations and 16 subsets with a 3 mm FWHM ramp filter.

### Multimodal image processing in neocortical and hippocampal regions

#### a) Generation of neocortical surfaces

To generate models of the cortical surface and measure cortical thickness, native T1w images of each participant were processed using FreeSurfer 6.0 (http://surfer.nmr.mgh.harvard.edu). Previous work has cross-validated this pipeline with histological analysis (Cardinale et al., 2014; Rosas et al., 2002) and manual measurements (Kuperberg et al., 2003). Processing steps have been described in detail elsewhere (Dale et al., 1999; Fischl et al., 1999). In short, the pipeline includes brain extraction, tissue segmentation, pial and white matter surface generation, and registration of individual cortical surfaces to the fsaverage surface template. The latter step aligns vertices among participants, whilst minimising metric distortion. Cortical thickness was calculated as the closest distance from the white matter to the pial boundary at each vertex. FreeSurfer quality control and manual edits were carried out by a single rater (AL) and included pial corrections and addition of control points, followed by reprocessing.

#### b) Hippocampal subfield surface mapping

We applied a recently developed approach for hippocampal subfield segmentation, the generation of surfaces running through the core of each subfield, and subsequent “unfolding” for surface-based analysis of hippocampal imaging features (Bernhardt et al., 2016; Caldairou et al., 2016; Kim et al., 2014; Vos de Wael et al., 2018). In brief, each participant’s native-space T1w image underwent automated correction for intensity non-uniformity, intensity standardization, and linear registration to the MNI152 template. Each image was processed using a multi-template surface-patch algorithm (Caldairou et al., 2016), which automatically segments the hippocampal formation into subiculum, CA1-3, and CA4-DG subfields. An open-access database of manual subfield segmentations and corresponding high-resolution 3T MRI data (Kulaga-Yoskovitz et al., 2015) was used to train the algorithm (https://www.nitrc.org/projects/mni-hisub25). The algorithm was previously validated in healthy individuals, where Dice coefficients above 0.8 were demonstrated across subfields even when millimetric T1w images were used as input (Caldairou et al 2016). Notably, the algorithm also generates medial sheet representations running through the core of each subfield, which allow for the sampling of intensity parameters with minimal partial volume effect, while guaranteeing point correspondence across subjects. Prior validation experiments in epileptic patients showed that these features reliably predict hippocampal histopathology and focus laterality (Bernhardt et al., 2017, 2016; Kim et al., 2014). After parametrizing subfield surfaces using spherical harmonic shape descriptors (Styner et al., 2006), a Hamilton-Jacobi approach generated a medial surface running through the central path of each subfield (Kim et al., 2014). To estimate local atrophy, we calculated columnar volume (Kim et al., 2014). This index is calculated as the distance between a vertex on the medial sheet of each subfield and the corresponding outer shell multiplied by the average surface area of the surrounding triangles between the subfield boundary and the medial surface. In prior work, we showed a high correlation between columnar volume and degrees of hippocampal cell loss in patients with temporal lobe epilepsy (Bernhardt et al., 2016).

#### c) PET-MRI integration

We mapped PET data to T1w imaging space generated by the pipelines in *a)* and *b)*, allowing for a surface-based integration of Aβ uptake with structural imaging features along neocortical and hippocampal subfield surfaces. In both cases, boundary-based procedures estimated the registration between a native PET image and the corresponding T1w images (Greve & Fischl, 2009), followed by vertex-wise interpolation within the cortical ribbon. Neocortical PET data were resampled to fsaverage5 to improve correspondence across measurements; in the case of hippocampal PET features, the sampling grid was already aligned via the spherical parameterization during processing (Kim et al., 2014). Following previous approaches (Rodrigue et al., 2012), neocortical and hippocampal Aβ values were normalized by mean cerebellar grey matter Aβ uptake, providing a standardized uptake value ratio (SUVR) per vertex. To control for cerebro-spinal fluid partial volume effects (CSF-PVE), each participant’s T1w image was skull stripped and segmented into tissue types and partial volume estimates (Zhang et al., 2001). CSF-PVE maps were mapped to neocortical and hippocampal surfaces. Using surface wide linear models, we controlled Aβ SUVR values for effects of CSF-PVE at each vertex in all participants.

### Quality Control and Final Sample Selection

The DLBS open-access data set contains structural and functional data with variable image quality. All T1w images, segmentations, and co-registrations were visually inspected by a single researcher (AL). We removed datasets with artefacts leading to inaccurate cortical segmentations (n=25) Additionally, data with inaccurate hippocampal segmentations (n=17), characterised by gross errors or inclusion of neighbouring white matter voxels, were removed. Following quality control, the final sample included 102 healthy individuals (69 females, 30-89 years, mean±SD age=59±16.1 years, 90.2% White/Caucasian), who were highly educated (mean±SD=16.1±2.2 years of education), and scored highly on the MMSE (mean±SD=28±1.1). The age of our sample was approximately normally distributed (**Supplementary Figure 1**).

### Statistical Analysis

Analyses were performed using SurfStat (Worsley et al., 2009) for Matlab (The Mathworks, Natick, MA, R2017B). All linear models outlined below additionally controlled for sex and years of education.

#### a) Regional analysis along neocortical and hippocampal surfaces

Surface-wide linear models examined effects of age on cortical thickness.

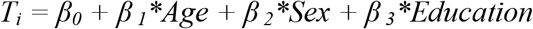

Where *T*_*i*_ is the thickness at vertex *i*, and *Age*, *Sex* and *Education* are the model terms and the *betas* the estimated model parameters. Similar models assessed the relationship between cortical Aβ deposition and age. We corrected for multiple comparisons using the false discovery rate (FDR) procedure (Benjamini & Hochberg, 1995). We selected a two-tailed p_FDR_ of <0.05. An analogous approach assessed age effects on columnar volume and Aβ across hippocampal subfield surfaces.

#### b) Relation to cognitive factors

We computed Pearson’s correlation coefficients and tested for associations between age and neuropsychological factor scores F1 and F2. Furthermore, we correlated mean cortical thickness, hippocampal volume, and Aβ values within significant clusters computed in *a)* with the factor scores. As for the previous analysis, findings were corrected using an FDR procedure.

#### c) Multimodal profiling based on connectome topographic mapping

We used previously derived maps of neocortical and hippocampal resting-state functional connectome gradients (Margulies et al., 2016; Vos de Wael et al., 2018) based on the human connectome project dataset (Van Essen et al., 2012) to stratify findings. In the neocortex, the first principal gradient describes a continuous transition from unimodal sensory areas via task-positive networks, such as the salience, dorsal attention, and fronto-parietal network, towards the DMN core regions (Margulies et al., 2016), thus recapitulating earlier models of a cortical hierarchy with low-level sensorimotor regions on the one end, and transmodal regions participating in higher-order functions on the other end (Mesulam, 1998). In the hippocampus, the first principal gradient of hippocampal connectivity runs from anterior to posterior regions across all subfields, with the former being more strongly connected to transmodal DMN than the posterior part (Vos de Wael et al., 2018). Both gradients were discretized into 20 bins, following a recent approach to stratify task-based fMRI data using connectome topographies (Murphy et al., 2018). Each bin contained the same number of vertices to ensure comparable sensitivity. We employed the same analysis as in *a)* and mapped significant t-values to the discretized gradient using a sliding window approach. Thus, each gradient bin had a specific t-value representing the effect of age on brain markers in that bin. This was plotted graphically against bin ordering (1-20). A linear model between bin ordering and the t-value within each bin explored interactions between gradient ordering and age effects on brain markers. To assess the relationship between gradient values and cognition, we computed the mean cortical thickness and amyloid values in each of the 20 gradient bins controlling for gender and level of education. The residual bin values were then fed into a linear model with factor score 1 and 2 and resultant t-values were plotted against bin ordering to produce gradient-cognition plots. A linear model between bin ordering and the t-values then explored interaction between gradient ordering and cognition. This approach, thus, provided a low-dimensional representation of structural and Aβ changes along the neocortical and hippocampal functional topography.

### Data and code availability

All data are based on the openly-shared Dallas Lifespan Brain Study dataset, available under http://fcon_1000.projects.nitrc.org/indi/retro/dlbs.html. Preprocessed and quality controlled surface feature data are available upon request. Surface-wide statistical comparisons were carried out using SurfStat for Matlab (http://www.math.mcgill.ca/keith/surfstat/). Gradient mapping tools used in this work are available via https://github.com/MICA-MNI/micaopen/diffusion_map_embedding.

## III. Results

### Effects of age on neocortical and hippocampal subfield markers

#### a) Morphology

Vertex-wise analysis revealed widespread reductions in cortical thickness with increasing age (FDR-corrected p-value, *p*_*FDR*_<0.025; Figure 1A). Laterally, clusters occupied bilateral frontal, central, temporal, and parietal cortices with a relative sparing of the orbitofrontal and occipital cortices. Medially, clusters occupied bilateral precuneus, cingulate, paracentral, superior frontal, fusiform and parahippocampal cortices. Considering the hippocampus, subfield analysis of columnar volume revealed effects predominantly in posterior portions, spanning subiculum, CA1-3, and CA4-DG bilaterally. Additional clusters were also observed in bilateral anterior CA1-3 (*p*_*FDR*_<0.025; FIGURE 1A).

**FIGURE 1.**
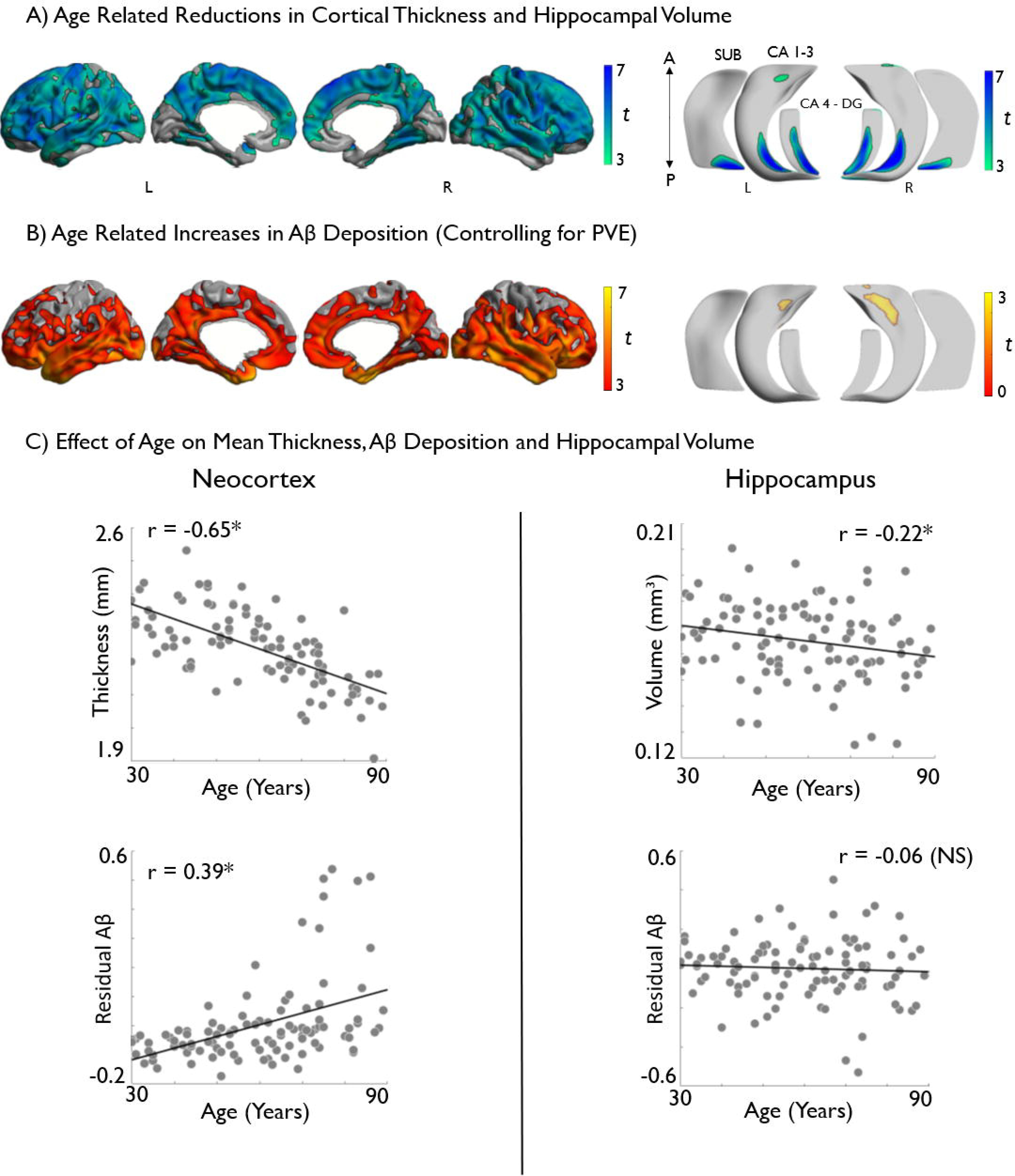
Analysis of grey matter morphology and Aβ deposition (normalized by cerebellar grey matter and controlled for CSF PVE) along neocortical (*left*) and hippocampal subfield (*right*) surfaces. Effects of age on **A)** cortical thickness and hippocampal volume across all subfields and **B)** on Aβ deposition. Models controlled for sex and education. Age related increases are shown in warm and decreases in cold colors. Regions significant at a two-tailed p_FDR_<0.05 are shown with black outlines, uncorrected trends relating to increased hippocampal Aβ are shown in semi-transparent **(B***, bottom right*). Correlations between age and markers of brain-aging are displayed in **C). \***Denotes statistical significance. *PVE = Partial Volume Effect. NS = Non-Significant*.

#### b) Aβ

We observed higher Aβ deposition with increasing age (*p*_*FDR*_<0.025) in a different spatial pattern than the cortical thickness findings. Specifically, we observed bilateral increases in predominantly limbic and transmodal cortices, encompassing lateral and medial temporal, insula, orbitofrontal, cingulate and midline parietal, as well as supramarginal regions, with a relative sparing of primary motor, occipital and mesial frontal cortices (Figure 1B). In the hippocampus, neither increases nor decreases in Aβ survived FDR-correction. At uncorrected thresholds (*p*<0.025), we observed mainly trends for increased Aβ in bilateral anterior CA1-3 (Figure 1B).

### Effects of age and imaging markers on cognition

Older age correlated with lower cognitive factors scores *i.e*., poorer fluid intelligence (F1: r=- 0.66, *p*<0.001) and episodic memory (F2: r=-0.48, *p*<0.001; Figure 2A).

**FIGURE 2.**
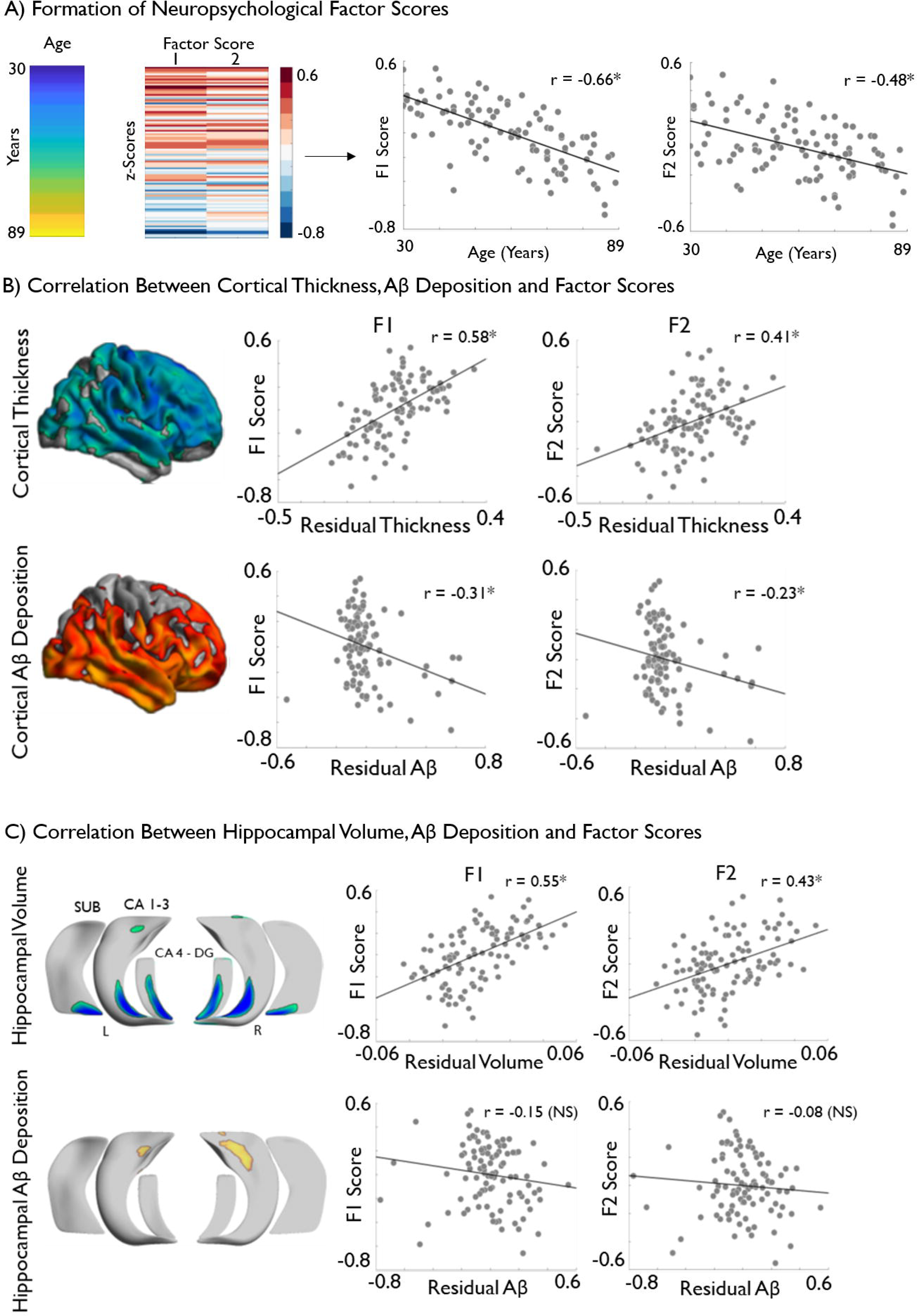
Associations between age-related brain markers and cognitive performance. **A)** Age of sample is displayed next to the results of the maximum likelihood common factor analysis with varimax rotation which identified two latent factors pertaining to measures of fluid intelligence (F1) and episodic memory (F2), respectively. The factor score matrix has been age-ordered with red indicating higher scores and blue indicated lower scores. Significant negative correlations between age and F1 and F2 scores are also displayed. **B)** *Post-hoc* correlation analysis, based on significant clusters of age-related cortical thickness and cortical amyloid deposition (See *Figure 1*) with F1 and F2. **C)** Correlation analysis between hippocampal volume and amyloid deposition with F1 and F2. Brain measures have been corrected for sex and education. Please see Figure 1 for details on the multiple comparison’s correction. *Denotes statistical significance. *SUVR = Standardized Uptake Value Ratio. NS = Non-Significant*

*Post-hoc* analysis between mean cortical thickness in regions of significant age effects (see *Figure 1A*) showed positive correlations with both F1 (r=0.58, *p*<0.001) and F2 (r=0.41, *p*<0.001), indicating better performance in individuals with higher thickness (Figure 2B). On the other hand, increases in mean Aβ deposition (see *Figure 1B*) related to lower scores on both F1 (r=-0.31, *p*<0.001) and F2 (r=-0.23, *p*<0.05) (Figure 2B). At the level of the hippocampus, we observed positive correlations between columnar volumes in regions of age effects and F1 (r=0.55, *p*<0.001) and F2 (r=0.43, *p*<0.001). Regarding hippocampal Aβ deposition in regions of uncorrected age effects, no significant correlations were observed with F1 (r=-0.15, *p*=0.13) nor with F2 (r=-0.08, *p*=0.3).

A *post-hoc* mediation analysis between age, cognition and brain markers in regions of significant age effects was performed. The brain markers selected as potential mediators were those which demonstrated a significant correlation with both factor scores (Figure 2B/C). As such, hippocampal Aβ deposition was not included. Following the methodology described by Zhao et al. (2010), we found cortical thickness (F1: *a*b* = 0.007 [CI = 0.005 – 0.009], F2: *a*b* = 0.004 [CI = 0.003 – 0.006]) cortical Aβ deposition (F1: *a*b* =-0.002 [CI = −0.004 - −0.002], F2: *a*b* = −0.001 [CI = −0.003 - 0.000]) and hippocampal volume (F1: *a*b* = 0.006 [CI = 0.005 – 0.008]. F2: *a*b* = 0.005 [CI = 0.003 – 0.006]) all to be significant mediators of the relationship between age and both F1 and F2. All brain markers were categorized as ‘Complimentary’ mediators (Zhao et al., 2010), indicating the likely presence of additional mediating variables on the relationship between age and cognition.

### Profiling of age effects on brain markers and cognition via connectome gradients

#### a) Age effects

Age-related cortical thinning was diffuse across the entire cortical functional gradient (*p*_*FDR*_ <0.025), with no marked difference between uni- and transmodal areas (t=-1.64, *p*=0.11) (Figure 3A). On the other hand, although age-related increases in Aβ deposition also occurred across the entire neocortical gradient, we observed a significant incline towards transmodal regions (t=6.96, *p*<0.001) (Figure 3A). Hippocampal age-effects were not as strong as in the neocortex and did not reach significance. Yet, effect sizes for age-related volume loss were larger towards the posterior aspect of the hippocampal ‘long-axis’ gradient (t=-9.51, *p*<0.001) while we observed trends for increased Aβ in anterior regions (Figure 3B).

**FIGURE 3.**
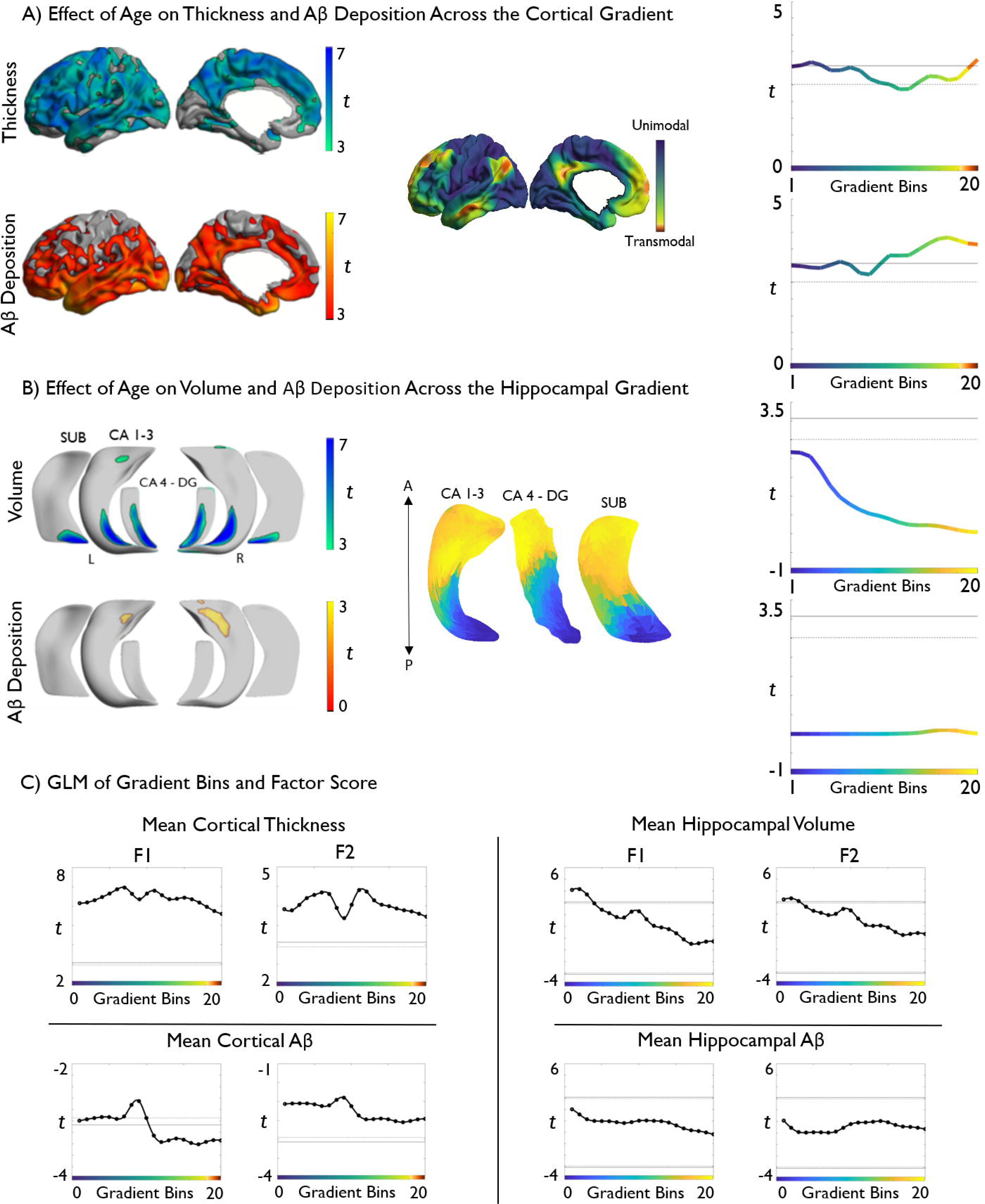
Topographic profiling of age effects and cognitive correlations in neocortical and hippocampal regions. **A)** Age effects on vertex-wise cortical thickness and Aβ deposition (*left*) were mapped to a reference space based on neocortical functional connectivity gradients (*centre*: adapted from Margulies et al., 2016) and an atlas of levels of laminar differentiation (*right*: adapted from Mesulam 1998; Paquola et al., 2018). Line profiles depict changes in t-statistics from primary sensory regions on the left side of the graph towards transmodal regions on the right *(centre)*. Boxplots show the median and range of t-statistics within levels of laminar differentiation *(right)*. Age-related effects on cortical thickness were consistently high across the entire neocortical gradient, and did not differ across levels of laminar differentiation. In contrast, Aβ shows higher effect of age towards the transmodal end of the gradient, corresponding to the paralimbic cortex. **B)** Vertex-wise age-effects on hippocampal subregion volume and Aβ deposition (*left*), mapped to a ‘long-axis’ reference space based on hippocampal functional connectivity gradients (*centre*, adapted from Vos de Wael et al., 2018), showing more elevated deposition in anterior subregions. Right hemisphere gradients were virtually identical. **C)** Gradient-based stratification of correlations between F1 and F2 on neocortical (*left panels*) and hippocampal measures (*right panels*). *Solid lines represent significant t-values using Bonferroni correction, whereas dashed lines represent significant t-values using FDR correction for multiple comparisons*.

#### b) Cognition

Considering cortical thickness, we found that measures across the entire neocortical gradient positively correlated with F1, with largest effects in sensory regions (t= −2.40, *p*<0.05), and with F2, which demonstrated no significant difference between sensory and higher order bins (t=-1.60, *p*=0.13). Aβ deposition across almost the entire gradient correlated with F1. However, in contrast to the thickness findings, we found that transmodal values were most closely related to F1 scores (t=-5.00, *p*<0.001). With regards to F2, although values across gradient bins did not reach significance, reduced scores on F2 were still observed as Aβ deposition increased in transmodal regions of the neocortical gradient (t=-6.80, *p*<0.001) (Figure 3C).

Hippocampal volume in low-level bins, corresponding to the posterior hippocampus correlated with higher F1 scores (t=-16.33, *p*<0.001) (Figure 3C) and higher F2 scores. Values in the posterior aspect of the gradient was again shown to have greater predictive power compared to the anterior portion (t=-15.44, *p*<0.001). Unlike the neocortical findings, gradient-wise hippocampal Aβ deposition did not correlate with scores for F1 and F2 (Figure 3C).

### Additional control analyses

Although main models included sex and education as control covariates, similar effects were observed using models that omitted their control (**Supplementary Figure 3**). Furthermore, we observed virtually identical results when additionally controlling for APOE-e4 genotype (**Supplementary Figure 4**), and when performing a subgroup analysis restricted to only non-APOE4-e4 carriers (**Supplementary Figure 5**).

Given that functional connectivity in healthy subjects has been shown to differ with age (Ferreira et al., 2016; Sala-Llonch et al., 2014), we also performed a separate control analysis in which we built functional connectivity gradients in the hippocampus and neocortex from a different healthy life span dataset (n=39, 20 females, age range: 18-77, mean±SD=45±22.9 years) (**Supplementary Methods**). Gradients estimated in this cohort were largely similar in overall shape to the original ones i.e. describing a system level transition from unimodal to transmodal areas in neocortices and the hippocampal long-axis. Gradient-stratified findings based on the lifespan functional data were thus virtually identical to the original findings based on the HCP cohort, both for the neocortex (**Supplementary Figure 6**) and hippocampus (**Supplementary Figure 7**).

Finally, we also mapped levels of laminar differentiation to our cortical surface models similar to our previous work integrating 3D histology and neuroimaging (Paquola et al. 2018). To this end, each cortical node was assigned to one of four levels of laminar differentiation (*i.e*., idiotypic, unimodal, heteromodal or paralimbic) derived from the seminal model of the cortical hierarchy formulated by Mesulam, which was built on the integration of neuroanatomical, electrophysiological, and behavioural studies in human and non-human primates (Mesulam, 1998). Laminar differentiation-based analysis confirmed highest aging related Aβ deposition in paralimbic transmodal areas (t>3.9), with effect sizes descending down the hierarchy i.e. heteromodal association areas and unimodal association areas (t=3.5) followed by idiotypic sensory and motor cortices (t=2.77). For thickness, findings were more diffuse as for the gradient based profiling with highest negative aging effects seen in unimodal and heteromodal association areas (t>3.3), followed by idiotypic (t=3.0), and then limbic areas (t=2.0).

## VI. Discussion

Our work targeted age-related differences in morphology and amyloid deposition across neocortical and hippocampal subregions and confirms widespread structural-metabolic differences with advancing age. Age effects on thickness were diffuse along the entire neocortical functional gradient, whereas effects on volume were stronger in posterior segments of the hippocampal long-axis. Regarding Aβ deposition, age-related increases were observed along the entire neocortical gradient with significantly stronger effects observed in higher-order transmodal neocortices while no gradient-based modulation of age effects was observed for hippocampal Aβ. Structural and amyloid measures correlated with behavioural indices of fluid intelligence and episodic memory, again in a topography-dependent manner, emphasizing the power of our analytical framework to compactly represent and conceptualize brain aging in the context of macroscale functional systems.

At a whole-brain level, our image processing strategy incorporated several desirable elements to combine imaging metrics of neocortical and hippocampal subregions. Indeed, unconstrained surface-based morphometric MRI and Aβ-PET analysis in neocortical regions extends work focussing on *a-priori* defined region-of-interest (Rodrigue et al., 2012; Thambisetty et al., 2010). Likewise, by unfolding its complex and interlocked anatomical organization, we could address subregional changes in the hippocampal formation along its long-axis, building upon prior work operating at the whole-hippocampus (Fjell et al., 2009) or subfield-wise level (de Flores et al., 2015). Notably, although Aβ sampling was carried out within the cortical ribbon to minimize cerebrospinal fluid partial voluming, we additionally controlled for these effects at each vertex using statistical techniques and normalized Aβ uptake data against cerebellar grey-matter values, a reference region thought to be relatively unaffected by aging (Rodrigue et al., 2012; Vandenberghe et al., 2010). These steps likely increased specificity, while minimizing morphological confounds. At the level of the neocortical surface, we could observe a divergence between the effects of age on thickness and Aβ. Indeed, while the former affected a widespread fronto-centro-temporal territory, in line with prior work (Fjell et al., 2009; McGinnis et al., 2011; Salat et al., 2004; Yang et al., 2016; Yao et al., 2012), increased Aβ was observed in a more restrictive and predominantly limbic-transmodal circuitry (Rodrigue et al., 2012; Sperling et al., 2009). Following FreeSurfer edits and quality control, results remained largely the same, supporting that little or no age-related differences reflect a genuine sparing with increasing age. The regional divergence of morphological and Aβ effects was paralleled in the hippocampus, where we observed age-related reductions in local columnar volume mainly in posterior segments, while Aβ was marginally increased in the proximity of the hippocampal head.

To conceptualize these spatial patterns in a framework that more closely relates to macroscale functional organization, we utilized novel topographic mapping techniques guided by resting-state functional connectivity information from a large sample of healthy adults. Specifically, we remapped cortical and hippocampal morphometric and Aβ measures according to the main axes of neocortical and hippocampal connectivity. Prior work has shown that neocortical connectivity variations follow a gradient running from unimodal towards transmodal regions while hippocampal connectivity gradually shifts along its long-axis (Margulies et al., 2016; Vos de Wael et al., 2018; Hong et al., 2019; Plachti et al., 2019). Representing neuroimaging data in this compact, and presumably hierarchical (Mesulam, 1998), reference frame can be seen as complementary to parcellation-based methods as it does not assume clear-cut boundaries between functional systems but rather gradual transitions when going from one network to the next. In keeping with our regional findings, topography-stratified analysis supported a difference between structural and Aβ changes relative to the neocortical axes. While age-effects on thickness were seen along the entire gradient, positive age-Aβ correlations were significantly larger towards the transmodal anchor. Our findings with respect to functional connectivity gradients is compatible with earlier work demonstrating age-related cortical thinning across multiple functional networks (Bajaj et al., 2017), and a selective vulnerability of higher-order midline networks to Aβ pathology (Mutlu et al., 2017; Myers et al., 2014; Palmqvist et al., 2017; Sperling et al., 2009). From a theoretical perspective, our approach complements earlier models assuming a spatial patterning of brain aging, for example models presuming that posterior structural and associated functional compromise may lead to increased activity in anterior regions (Davis et al., 2008; Grady et al., 1994; Payer et al., 2006), or even more general accounts that assume the engagement of supplementary networks to preserve cognitive function in the face of diffuse neurofunctional decline (Park & Reuter-Lorenz, 2009; Reuter-Lorenz & Park, 2014). Structural decline of neurotransmitter systems throughout the brain may also result in functional changes, including attenuation of neuronal gain control, resulting in suboptimal cognition (Li et al., 2001; Li & Rieckmann, 2014). In fact, our results provide support for hierarchy-specific shifts, whereby diffuse structural changes result in compensatory recruitment of higher-level regions. In other words, cortical atrophy across a large territory could evoke increased functional demands on higher-order default mode and frontal-parietal networks. Though not fully understood, the increase in activity, connectivity, and metabolic stress may in turn increase the susceptibility of higher-order networks to Aβ pathology (Bero et al., 2011; Buckner et al., 2005; Lehmann et al., 2013). Regarding the hippocampal gradient, we observed larger age-related volume loss in posterior aspects of the hippocampal long-axis, supporting earlier literature demonstrating a vulnerability of the posterior hippocampus in aging (Pruessner, Collins, Pruessner, & Evans, 2001). We found no significant effect of age on hippocampal Aβ deposition across the entire hippocampal gradient, which is in keeping with our regional findings demonstrating only trends for increased deposition in the hippocampal head. We believe our lack of significant findings relating to Aβ across the hippocampus could be the result of low sensitivity of PET imaging to Aβ deposition in this structure. The unique anatomy of the hippocampal formation, coupled with its predisposition towards partial volume errors, has been hypothesised to reduce the reliability of PET imaging in this region (Sabri, Seibyl, Rowe, & Barthel, 2015). Nevertheless, we deemed it worthwhile to explore hippocampal Aβ, given theoretical benefits when studying metabolic data with subfield-surface analytics that offer reduced partial volume effect during parameter sampling and high spatial specificity.

With regards to the cognitive substrates of our findings, thickness reductions across the neocortical gradient related to lower scores on measures of episodic memory and fluid intelligence, supporting a contribution of whole-cortex morphological integrity to this faculty (Fjell et al., 2006; Schretlen et al., 2000). As effect sizes were somewhat higher in unimodal portions of the gradient, our data may underline the contribution of externally-oriented attention networks to fluid intelligence (Majerus et al., 2012), with particularly the dorsal attention network being proximal to sensory and sensorimotor anchors on the neocortical gradient (Margulies et al., 2016). The dorsal attentional network and sensorimotor regions are densely interconnected, and previous research has shown that externally-oriented operations broadly contribute to fluid intelligence (Hearne et al., 2016). With respect to Aβ, particularly transmodal neocortical increases related to worse scores, with effects significant for fluid intelligence but only trending for memory-related factor scores. Although not explored here, the observed structural and metabolic change across neocortical and hippocampal regions may reflect disruptions of large-scale structural networks, negatively affecting cognitive functions. Previous work indeed showed a decline white matter network efficiency with age (Collin & Van Den Heuvel, 2013; T. Zhao et al., 2015), with long-range connections and higher-order cognitive networks demonstrating considerable vulnerability (Montembeault et al., 2012; Spreng & Turner, 2013; Tomasi & Volkow, 2012).

When controlling for APOEe4 genotype status, we observed no modulation or differences in cognitive score. Given recent data indicative of a significant effect of genotype status on cognition in aging (Schiepers et al., 2012), this result was quite surprising. One potential explanation for this finding is that the relatively small sample of APOEe4 carriers (n=23) in this current study was not large enough to demonstrate any significant effect. However, our results do lend support to earlier work demonstrating no effect of APOEe4 status on cognition in healthy aging (Mayeux, Small, Tang, Tycko, & Stern, 2001; Pendleton et al., 2002; B. J. Small et al., 2000; Brent J. Small, Basun, & Bäckman, 1998; Smith et al., 1998). Furthermore, in studies that do find an effect of APOEe4 on cognition, the effect is not consistent across cognitive domains, with attention, primary memory, verbal ability, visuospatial skill and perceptual speed demonstrating no significant deficits as a result of genotype status (Brent J. Small, Rosnick, Fratiglioni, & Bäckman, 2004; Wisdom, Callahan, & Hawkins, 2011).

A potential limitation to this study is that we were unable to control for subjective cognitive decline (SCD). SCD was assessed in the DLBS using the Metamemory in Adulthood questionnaire (Chen, Farrell, Moore, & Park, 2019), however, this data has yet to be released. SCD has been related to mesiotemporal atrophy and functional connectivity changes (Fan et al., 2018; Verfaillie et al., 2018), and might thus have been an interesting variable to relate to neocortical and hippocampal measures in the newly proposed gradient space. Furthermore, the DLBS only screened against physical health, neurological health, and MMSE>26 to cover a large range of individuals falling under a healthy aging umbrella. Measures of preclinical Alzheimer’s disease, including Amyloid beta SUVR cut-off points and/or Scheltens visual rating scale for mesiotemporal atrophy (Philip Scheltens, Launer, Barkhof, Weinstein, & van Gool, 1995) were thus not used for subject exclusion. It is, therefore, possible that some participants might have suffered from preclinical stages of Alzheimer’s disease with still high MMSE scores. Finally, please note that we restricted our analysis to 102/144 cases with higher-quality MRI data. Although this nominally reduced statistical power, focussing on cases with high-quality imaging data and quality controlled hippocampal and cortical segmentations may improve inferences. In fact, it might even be more sensitive than an assessment of a larger dataset with potential confounds in image quality and surface extractions.

In conclusion, our work presents a novel approach to represent age-related differences in brain structure, metabolism, and cognition. In addition to supporting previous work indicative of structural and metabolic change in the aging brain, the use of a compact analytical framework to relate brain-based biomarkers to macroscale functional systems allowed for novel insights into the interplay between pathological deposits and structural compromise, and how this subsequently impacts upon cognition in the healthy aging population.

## Acknowledgements

We like to thank the investigators of the DLBS and associated funding sources for making their data available, and INDI/FCP1000 for hosting the imaging data. Dr Casey Paquola was supported by the Transforming Autism Care Consortium. Reinder Vos de Wael MSc was supported by the Savoy Foundation for Epilepsy Research. Sara Larivière MSc was supported by a FRQS Fellowship. Drs Neda and Andrea Bernasconi were funded by the Canadian Institutes of Health Research (CIHR) and received salary support from the Fonds de la Recherche du Quebec – Santé (FRQS). Dr Boris Bernhardt acknowledges research support from the National Science and Engineering Research Council of Canada (NSERC Discovery-1304413), the Canadian Institutes of Health Research (CIHR FDN-154298), SickKids Foundation (NI17-039), as well as salary support from the Fonds de la Recherche du Quebec - Santé (FRQS Junior 1 Research Scholar).

