## Supplementary Materials for "Targeting Age-Related Differences in Brain and Cognition with Multimodal Imaging and Connectome Topography Profiling"

*Lowe et al.*

### SUPPLEMENTARY METHODS

#### Participants

20 young (mean age±SD=24±4 years, 10 females) and 19 older (mean age±SD=68±5 years, 10 females) adults were selected from a larger ongoing study on heathy aging at Cornell University. Participants were recruited from the community and completed a comprehensive cognitive test battery and magnetic resonance image (MRI). Young and old subjects were combined to create a final sample of 39 subjects (20 females, age range: 18-77 years, mean±SD=45±22.9 years) from which the neocortical and hippocampal connectivity gradients were generated. To be eligible for the study, participants had to be between the ages of 18-35 (Young) or over age 60 (Old). Exclusion criteria included any MRI contraindications and/or a history of neurological, neuropsychiatric, or cardiovascular disease. All participants were cognitively normal based on self-report on intake and cognitive screen (MMSE > 26). All participants provided informed consent consistent with procedures approved by the Institutional Review Board of Cornell University.

#### Structural MRI Acquisition

Imaging data for participants recruited at Cornell University were acquired using 3T GE Discovery MR750 scanner (General Electric, Milwaukee, United States) with a 32-channel receive-only phased-array head coil at the Cornell Magnetic Resonance Imaging Facility in Ithaca. Anatomical scans were acquired with a T1-weighted volumetric MRI magnetization prepared rapid gradient echo (repetition time (TR)=2530ms; echo time (TE)=3.44ms; flip angle (FA)=7°; 1.0mm isotropic voxels, 176 slices). Anatomical scans were acquired during one 5m25s run with 2x acceleration with sensitivity encoding. Structural data was corrected for non-uniform intensities, affine-registered to Montreal-Neurological Institute (MNI) atlas and skull-stripped using FSL.

#### Multi-Echo Resting-State fMRI Acquisition

Participants completed one 10 min 6 second long resting-state multi-echo BOLD functional scan with eyes open, while blinking and breathing normally in the dimly lit scanner bay. Functional scans were acquired using a multi-echo echo-planar imaging (ME-EPI) sequence with online reconstruction (TR=3000ms; TE's=13.7, 30, 47ms; FA=83°; matrix size=72x72; field of view (FOV)=210mm; 46 axial slices; 3.0mm isotropic voxels].

#### Multi-Echo Resting-State fMRI Pre-Processing

Multi-echo fMRI facilitates removal of noise components from resting fMRI datasets (Kundu et al., 2013, 2012; Power et al., 2018). The method relies on the acquisition of multiple echoes, enabling direct measurement of T2* relaxation rates. Blood oxygen level dependent (BOLD) signal can then be distinguished from non-BOLD noise based on TE dependence. To remove noise components (*e.g.,* CSF, movement) from the data, independent components analysis is used to recombine and analyze the multiple echo-times. This method has shown to be successful in denoising BOLD signal of motion and physiological artifacts (Kundu et al., 2013, 2012). In the current work, data were preprocessed with multi-echo independent components analysis (ME-ICA) version 3.2 beta1 (https://github.com/ME-ICA/me-ica/blob/master/meica.py) and aligned to MNI space. ME-ICA processing was run with the following options: -e 13, 30, 46, -b 4v; –no_skullstrip; –space = young-old_n100_template.nii.gz. Anatomical images were skull stripped using FSL-BET prior to processing. Young-old_n100_template.nii.gz representes an averaged MNI152-space template of a larger sample of younger and older adults. Further filtering was omitted, as ME-ICA has shown to be successful in denoising BOLD signal of artifacts (Kundu et al., 2013, 2012). Components identified as signal were visually inspected for further quality control. All included participants had at least 10 signal components.

#### Generation of principle connectivity gradient in neocortex and hippocampus

*a) Neocortical.* To generate models of the cortical surface, each partipant's T1-weighted image was processed following the same methods outlined in the manuscript (see *Materials and Methods* at *P.6*). Resting-state fMRI time series were then mapped to surface-space, followed by resampling to fsaverage5. The principle gradient of neocortical connectivity was generated following the methodology outlined by Margulies et al. (2016) and Vos de Wael et al (2018). In brief, functional connectivity matrices were calculated from time-series cross-correlations, followed by Fisher r-to-z-transformation. Z-score matrices were averaged acoss all participants. For each region, we retained z-scores for the 10% strongest connections, with all others zeroed. An affinity matrix was subsequently generated that captures similarity in connectivity, based on a norm angle formulation. The principal gradient was then derived from this affinity matrix using diffusion map embedding (Coifman et al., 2005), which serves to recover a low dimensional embedding from high-dimensional connectivity data.

*b) Hippocampus.* To generate the principal gradient of hippocampal connectivity, we followed the methodology previously described by Vos de Wael et al. (2018). In brief, we first sampled each subject's volumetric rs-fMRI time series to the average medial sheet mesh across each subfield and calculated Pearson correlation coefficient maps across all vertices. Correlation coefficient maps were r-to-z converted followed by norm angle affinity matrix calculation and diffusion map embedding as in *a)*.

A MATLAB implementation of the diffusion embedding algorithm is available at https://github.com/MICA-MNI/micaopen/.

### SUPPLEMENTARY FIGURES

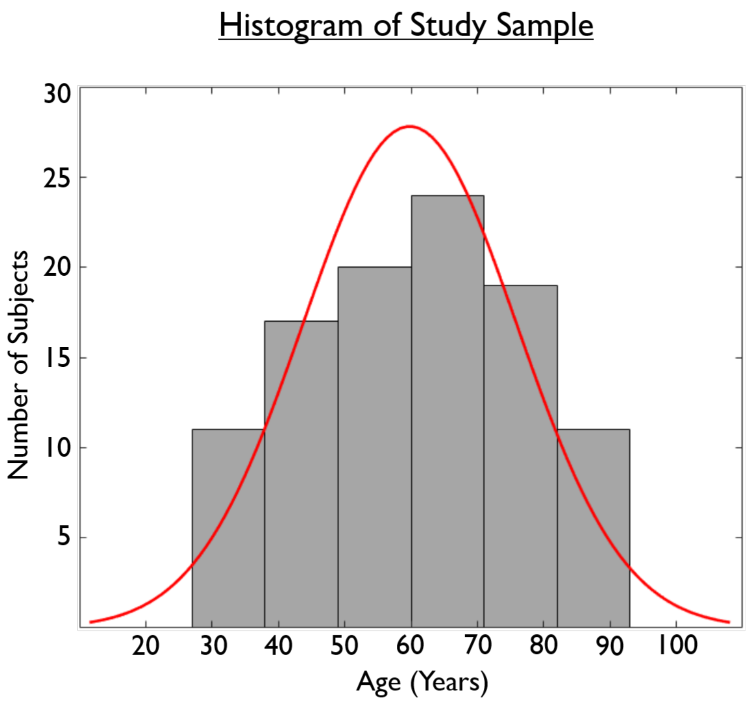


**SUPPLEMENTARY FIGURE 1.** Panel displays a histogram showing the final study sample (n=102) to be normally distributed.

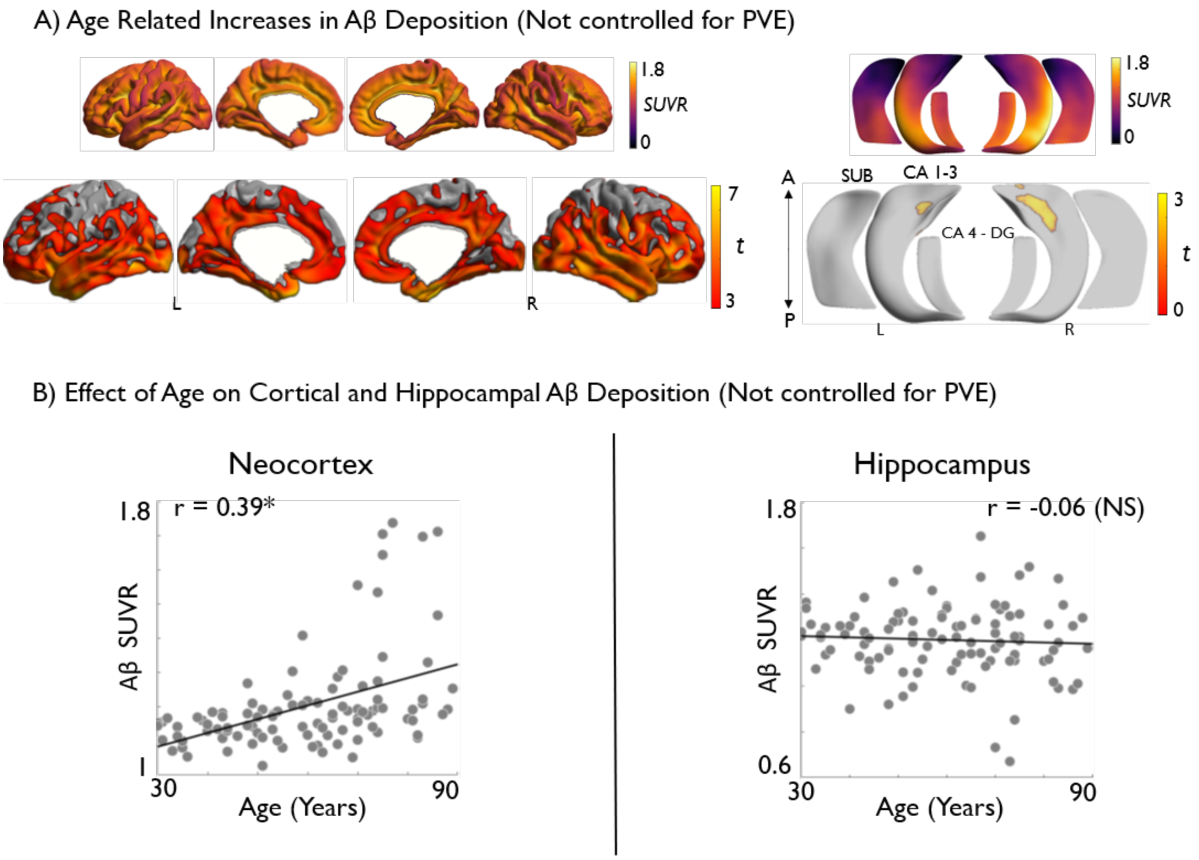


**SUPPLEMENTARY FIGURE 2.** Analysis of Aβ deposition along neocortical (*left*) and hippocampal subfield (*right*) surfaces. Aβ deposition is normalized by cerebellar grey matter but not controlled for PVE CSF. Effects of age on **A)** Aβ deposition. Models controlled for gender, age and education. A two-tailed pFDR of <0.05 was used. Age related increases are shown in warm colors. Regions significant at $p_{FDR}<0.05$ are shown with black outlines, uncorrected trends relating to increased hippocampal Aβ are shown in semi-transparent. Correlations between age and Aβ are displayed in **B).** *SUVR = Standardized Uptake Value Ratio. NS = Non-Significant.*

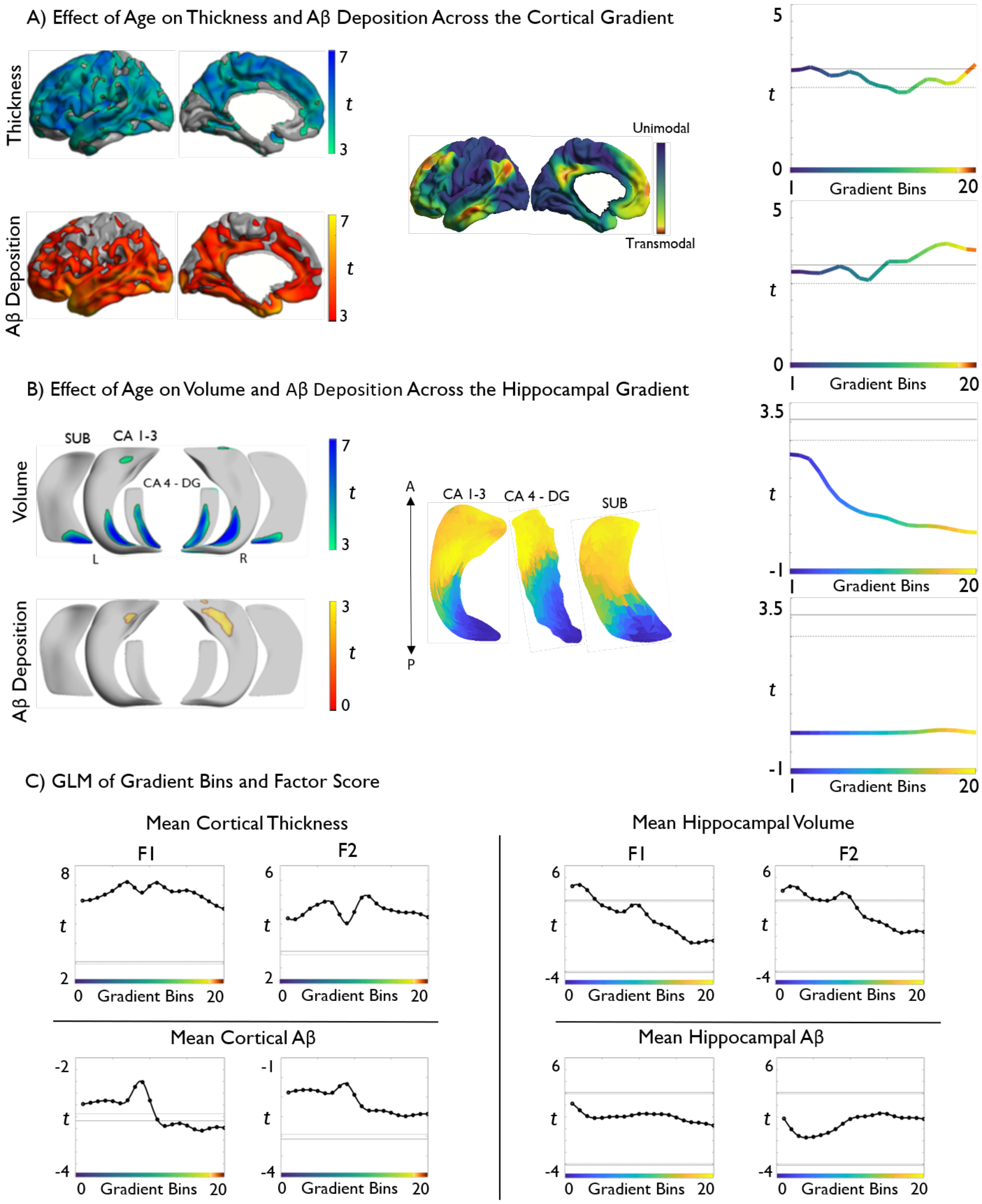


**SUPPLEMENTARY FIGURE 3.** Topographic profiling of age effects **(A, B)** and cognitive correlations **(C)** in neocortical and allocortical hippocampal regions**.** Models do not control for effect of gender or education.

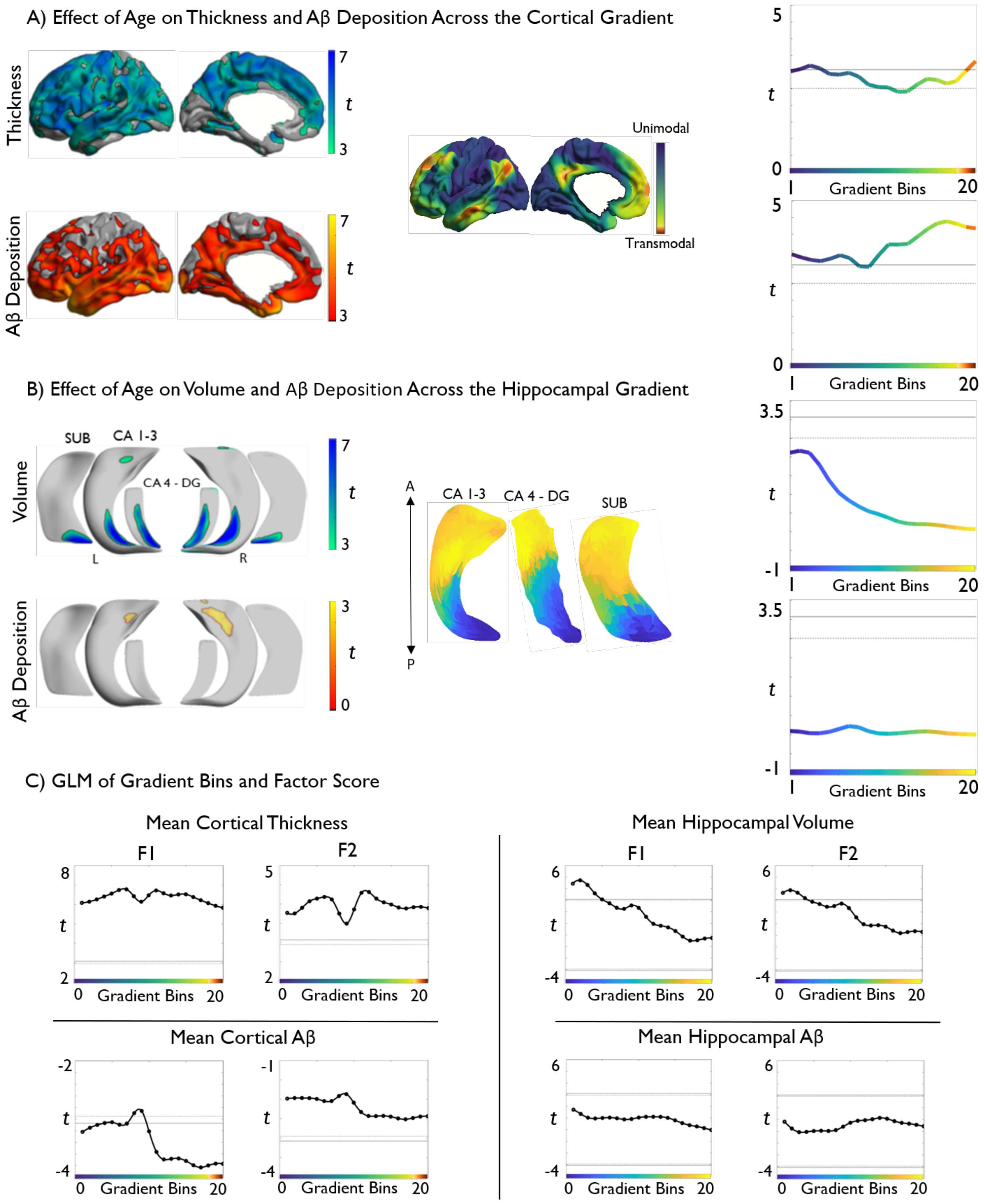


**SUPPLEMENTARY FIGURE 4.** Topographic profiling of age effects **(A, B)** and cognitive correlations **(C)** in neocortical and allocortical hippocampal regions**.** Models control for effect of gender, education and APOEe4 genotype status (n=99).

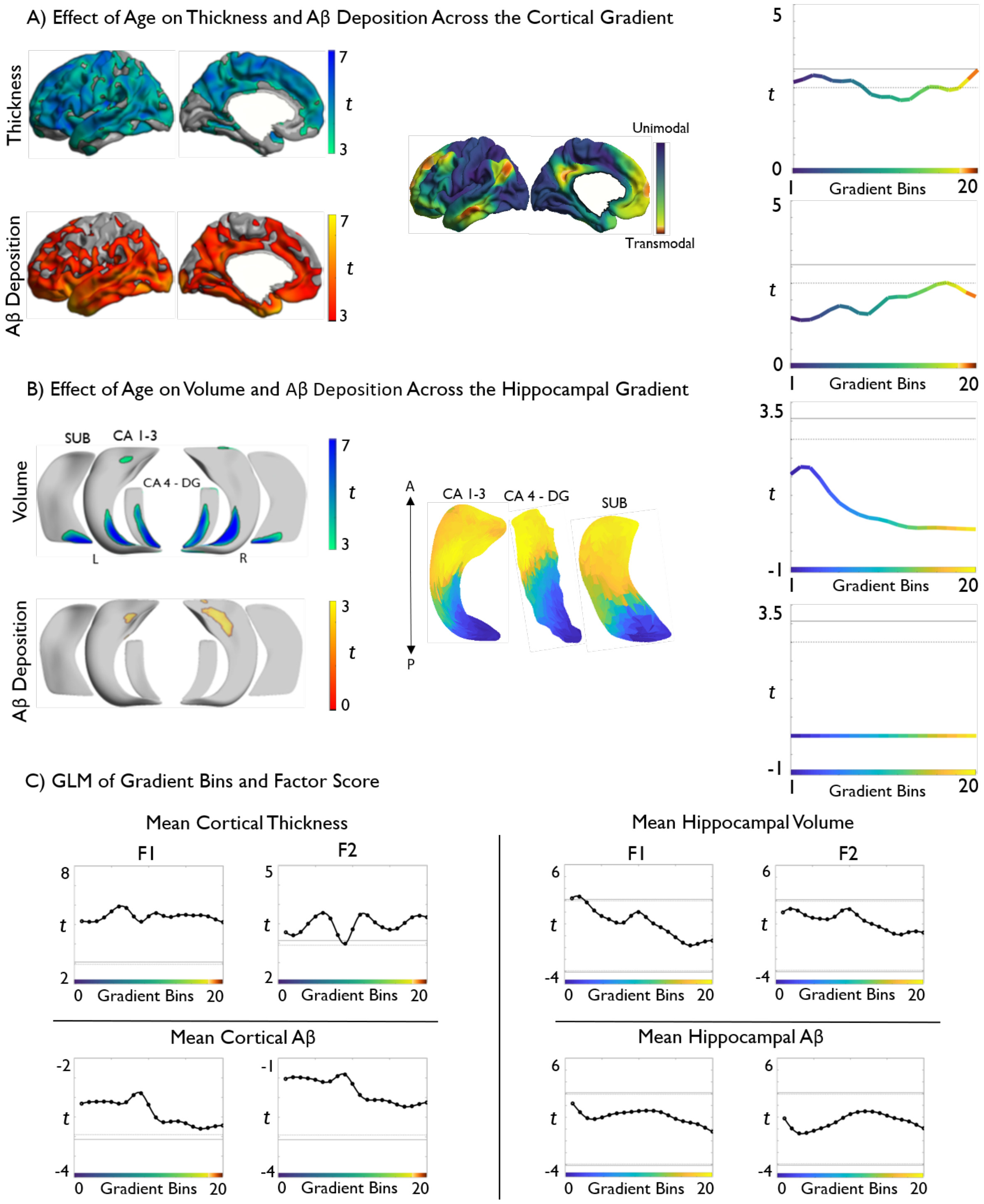


**Supplementary Figure 5.** Topographic profiling of age effects **(A-B)** and cognitive correlations **(C)** in neocortical and allocortical hippocampal regions. Subjects with e3/e4 and e4/e4 genotype were removed from analysis (n = 23). Models control for effect of gender and education.

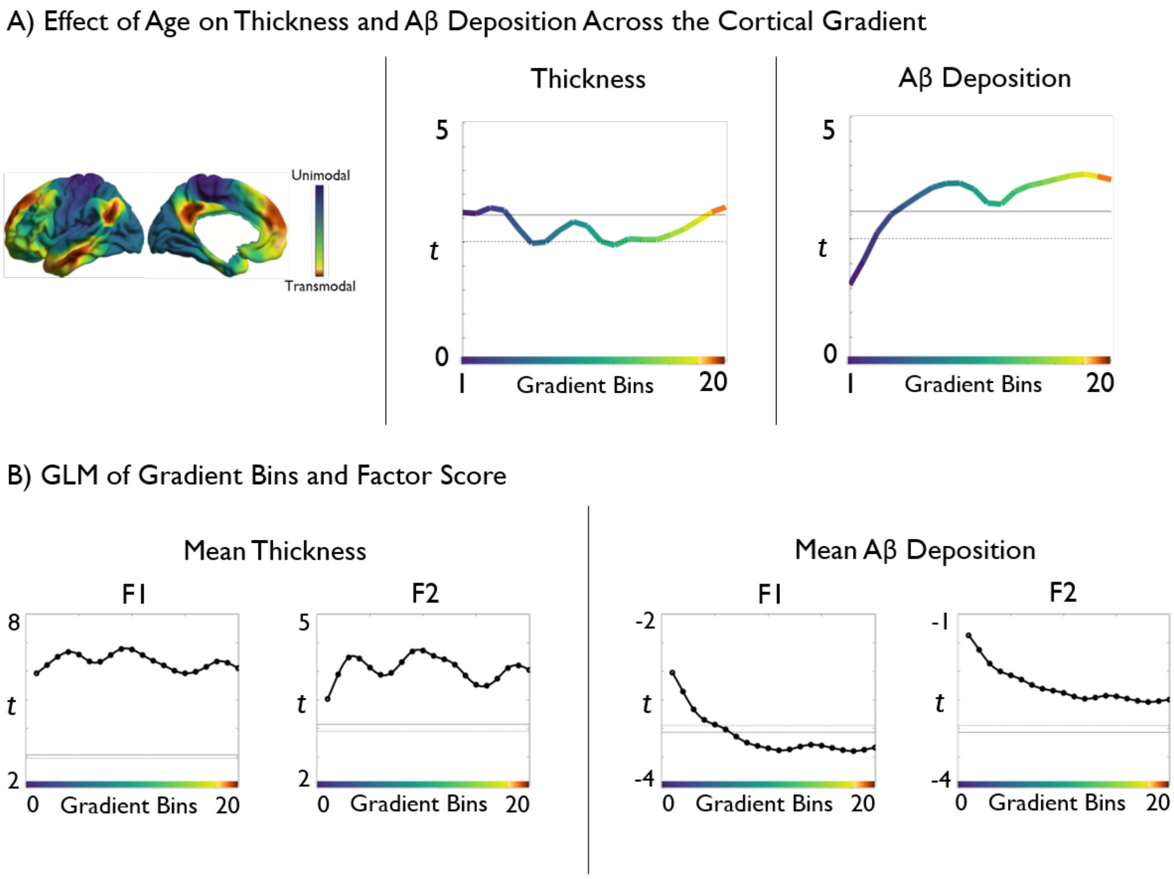


**SUPPLEMENTARY FIGURE 6.** Topographic profiling of age effects **(A)** and cognitive correlations **(B)** in neocortical regions. Functional connectivity gradient *(Top left panel)* was derived from a healthy lifespan data set (n=39, 20 females, age range: 18-77, mean±SD=45±22.9 years). Both age effects and cognitive correlations demonstrated similar results to those observed when using the original connectivity gradient derived from the HCP data set.

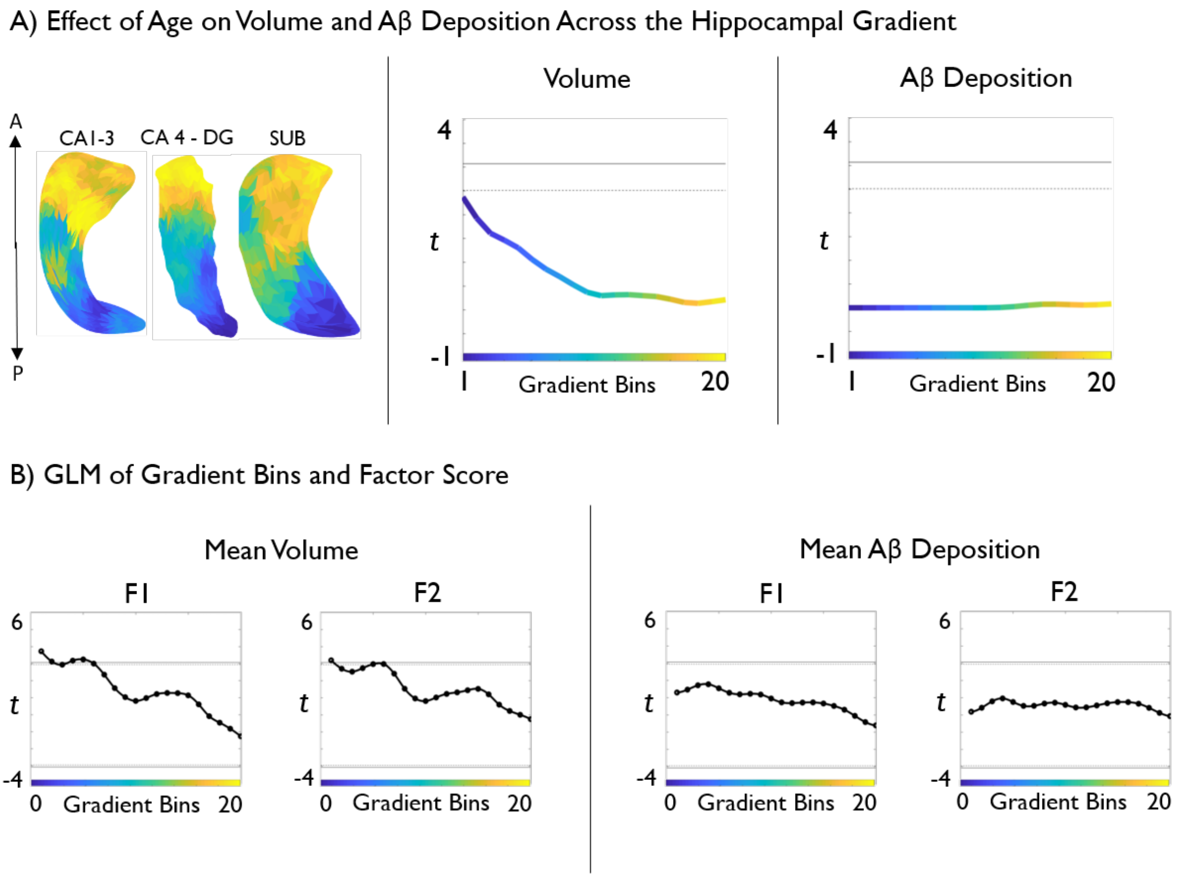


**SUPPLEMENTARY FIGURE 7.** Topographic profiling of age effects **(A)** and cognitive correlations **(B)** in hippocampal regions. Functional connectivity gradient *(Top left panel)* was derived from a healthy lifespan data set (n=37, 19 females, age range: 18-77, mean±SD=45±23.1 years). Both age effects and cognitive correlations demonstrated similar results to those observed when using the original connectivity gradient derived from the HCP data set.

### SUPPLEMENTARY TABLES

**SUPPLEMENTARY TABLE 1**. Factor Loading and tests**.**

| TEST | MEASURE | FACTOR 1 LOADING | FACTOR 2 LOADING |
|---|---|---|---|
| **MMSE** | General Cognition | 0.3309 | 0.2999 |
| **RAVENS PROGRESSIVE MATRICES** | Fluid Reasoning | 0.7903 | 0.2377 |
| **LETTER NUMBER SEQUENCING** | Working Memory | 0.3908 | 0.2290 |
| **HVL IMMEDIATE RECALL** | Episodic Memory | 0.1834 | 0.8257 |
| **HVL DELAYED RECALL** | Episodic Memory | 0.1275 | 0.8628 |
| **HVL RECOGNITION** | Episodic Memory | 0.1575 | 0.5421 |
| **ETS ADVANCE VOCABULARY** | Crystalized Ability | 0.0908 | 0.1975 |
| **ETS LETTER SETS** | Fluid Reasoning | 0.7412 | 0.1524 |
| **DIGIT SYMBOL** | Processing Speed | 0.6140 | 0.2558 |
| **DIGIT COMPARISON** | Processing Speed | 0.5333 | 0.3214 |
| **CANTAB VERBAL RECOGNITION** | Working Memory | 0.3844 | 0.4533 |
| **CANTAB SPATIAL WORKING** | Working Memory | 0.5666 | 0.0339 |
| **CANTAB STOP SIGNAL** | Executive Function | 0.2716 | 0.1745 |
| **CANTAB STOCKINGS OF CAMBRIDGE** | Executive Function | 0.5160 | 0.1476 |
